# Local interactions between steady-state visually evoked potentials at nearby flickering frequencies

**DOI:** 10.1101/2022.01.10.475604

**Authors:** Kumari Liza, Supratim Ray

## Abstract

Steady-state visually evoked potentials (SSVEP) are widely used to index top-down cognitive processing in human electroencephalogram (EEG) studies. Typically, two stimuli flickering at different temporal frequencies (TFs) are presented, each producing a distinct response in the EEG at its flicker frequency. However, how SSVEP responses in EEG are modulated in the presence of a competing flickering stimulus just due to sensory interactions is not well understood. We have previously shown in local field potentials (LFP) recorded from awake monkeys that when two overlapping full screen gratings are counter-phased at different TFs, there is an asymmetric SSVEP response suppression, with greater suppression from lower TFs, which further depends on the relative orientations of the gratings (stronger suppression and asymmetry for parallel compared to orthogonal gratings). Here, we first confirmed these effects in both male and female human EEG recordings. Then, we mapped the response suppression of one stimulus (target) by a competing stimulus (mask) over a much wider range than the previous study. Surprisingly, we found that the suppression was not stronger at low frequencies in general, but systematically varied depending on the target TF, indicating local interactions between the two competing stimuli. These results were confirmed in both human EEG and monkey LFP and electrocorticogram (ECoG) data. Our results show that sensory interactions between multiple SSVEPs are more complex than shown previously and are influenced by both local and global factors, underscoring the need to cautiously interpret the results of studies involving SSVEP paradigms.

## Introduction

Steady-state visually evoked potentials (SSVEPs) are stimulus-locked oscillatory electrical signals generated in response to a temporally periodic visual stimulus (Regan & D., 1989; Regant et al., 1988), with the response frequency strictly following the input stimulation frequency. SSVEP amplitude and phase are stable over time, have a high signal-to-noise ratio and are relatively immune to artefacts (Norcia et al., 2015; Vialatte et al., 2010). Apart from the response at the stimulation frequency, harmonic and intermodulation (IM) components are also obtained (Zemon & Ratliff, 1984), which could convey important information about the non-linear neural interactions in the visual system. In a non-invasive approach like electroencephalogram (EEG) that captures the synchronous activity of many neurons, it is often difficult to isolate the responses due to the presentation of multiple stimuli. By flickering the stimuli at different temporal frequencies (TFs), SSVEPs present a convenient way to segregate the stimulus-specific responses. Many cognitive paradigms such as visual attention (Andersen et al., 2012; Ding et al., 2006; Müller et al., 2006; Toffanin et al., 2009), binocular rivalry (Wang et al., 2004) and working memory (Silberstein et al., 2001) extensively use multiple stimuli tagged with distinct frequencies whilst SSVEP responses are measured. But the interactions of SSVEP responses due to multiple concurrently presented flickering stimuli have not been thoroughly investigated.

In masking studies, two stimuli are simultaneously presented in the same visual space leading to neural interactions between the competing stimuli (Legge & Foley, 1980). The response of the one grating, often called target grating, is attenuated due to another mask grating, even when the second grating fails to elicit any response when presented alone (Boynton & Foley, 1999; Candy et al., 2001; Foley, 1994; Morrone| et al., 1982; Tsai et al., 2012). Such effects have often been explained using a normalization model (Carandini et al., 1997; Carandini & Heeger, 1994, 2012; Heeger, 1992), and provide a framework to study interactions between two competing flickering stimuli.

We recently studied interactions between SSVEPs in local field potential (LFP) signals recorded from the primary visual cortex of awake macaques (Salelkar & Ray, 2020); the possible interactions and previous findings are shown in Figure 1. The SSVEP response or “gain” profile (Figure 1A) shows that stimuli flickering at ~10 Hz produced the strongest response. According to the normalization model, this TF should produce the strongest suppression to a competing (target) TF if the normalization strength only depended on SSVEP gain (Figure 1D; SSVEP gain-specific). By presenting the target at 16 Hz and masks at nearby frequencies (marked in magenta circles), we confirmed this asymmetric suppression in which lower mask TFs produced greater suppression (Figure 1D), but only when the constituent gratings were parallel (for orthogonal gratings, the suppression was non-specific as shown in Figure 1C). However, this can also be explained in a model in which normalization is stronger at lower frequencies (Figure 1E), potentially due to low-pass filtering in the normalization pool (Tsai et al., 2012). To distinguish these two possibilities, we presented the target at 8 Hz, for which the two hypotheses have opposite predictions, and found evidence in favour of Low-frequency suppression (Figure 1E).

**Figure 1:**
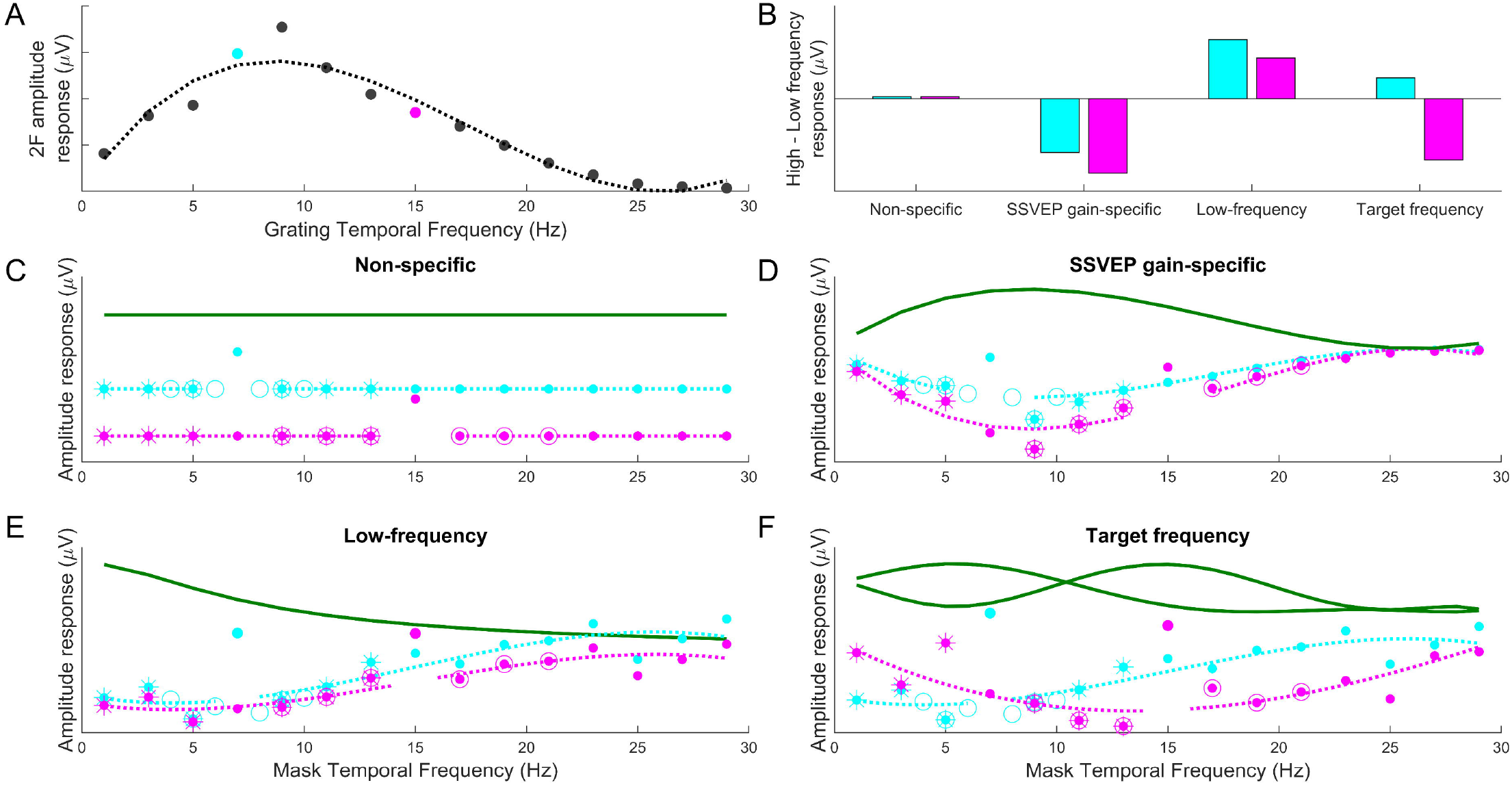
Different mechanisms of target and mask frequency interactions. **(A)** Averaged ECoG data from Monkey 1 when a single counterphase grating presented at 25% contrast and varying temporal frequency was used to obtain the SSVEP response function. Curve indicate a polynomial fit (degree = 3). Cyan and magenta dots represent the two target frequencies used in the main study **(B)** Difference between the average SSVEP amplitudes at 9, 11 and 13 Hz and 1, 3 and 5 Hz (marked as asterisks), under different hypotheses as discussed in C to F. Cyan and magenta bars correspond to the conditions when the target frequency was 7 and 15 Hz, respectively. **(C)** Non-specific interaction: Cyan and magenta curve depicts 7 and 15 Hz target frequency response as a function of temporal frequency of mask grating. The green curve is the underlying suppression function which is a horizontal line indicating the suppression of target SSVEP response is independent of the mask frequency. Solid dots are the different mask frequencies used in this study whereas the circled data are the frequencies used in our previous study (we had actually used 8 and 16 Hz target frequencies in the previous study, but shifted to 7 and 15 Hz to prevent subharmonic responses at 4 and 2 Hz). **(D)** SSVEP gain-specific interaction: The suppression function is same as the SSVEP response function indicating strongest suppression for the temporal frequency range that elicit maximum SSVEP response which is around 10 Hz for both the target frequencies (7 Hz: cyan curve; 15 Hz: magenta curve). **(E)** Low-frequency tuned interaction: The suppression signal (green curve) has more strength at progressively lower frequencies, potentially due to a low-pass filtering action on the normalization signal **(F)** Target frequency dependent interaction: The suppression function varies with the target frequency and is maximum in the vicinity of the target frequency.

In this study, we first tested whether these results also hold in human EEG recordings. Further, we characterized the response suppression profile by presenting masks over a much larger frequency range (1-29 Hz, dots in Figure 1). Surprisingly, the suppression profile was found to be gain-specific (Figure 1D), not low-frequency (Figure 1E), for the higher target frequency (magenta trace). Note that in all these models, the suppression has the same “global” profile (shown in green traces), irrespective of the target frequency. However, it is possible that the suppression profile itself depends on the target frequency, as shown in Figure 1F. By presenting the target at a lower frequency as well (blue trace), we found evidence in favor of this hypothesis, in both human EEG and monkey LFP and electrocorticogram recordings.

## Materials and Methods

### Human EEG recordings

Three experiments were conducted with 10, 10 and 11 subjects (total subjects: 31; 14 females and 17 males) recruited from the student community of Indian Institute of Science, Bangalore (mean age: ~23.1; range: 21-28 years). All experimental procedures were approved by the Institute Human Ethics Committee of Indian Institute of Science. Informed consent from all subjects was taken before the start of the experiment, and monetary compensation was provided for their voluntary participation. EEG signals were recorded from 8 active electrodes (actiCAP) using BrainAmp DC EEG acquisition system (Brain Products GmbH). The electrodes were placed in the parieto-occipital and occipital areas based on the international 10 – 10 system. The electrodes used were PO7, PO3, POz, PO4, PO8, O2, Oz and O1. Raw signals were filtered between 0.016 Hz (first-order filter) and 1000 Hz (fifth-order Butterworth filter), sampled at 2500 Hz and digitized at 16-bit resolution (0.1 μV/bit). Reference electrode was at FCz. The impedance was maintained below 20kΩ throughout the recording session.

### Animal recordings

Animal experiments were approved by the Institutional Animal Ethics Committee of the Indian Institute of Science and were conducted in accordance with the guidelines approved by the Committee for the Purpose of Control and Supervision of Experiments on Animals. Appropriate measures were taken during the experiment to minimize pain and discomfort to the animals. Two adult female bonnet monkeys *(Macaca radiata:* 3.3 and 3.4 kg) were used in this study. A titanium head post was implanted over the frontal region under general anaesthesia prior to training. Monkeys were then trained for passive visual fixation tasks. The first monkey was the same as Monkey 1 in our previous report on SSVEP interactions (Salelkar and Ray, 2020), but with a different “hybrid” array implanted on the other hemisphere that consisted of 81 (9×9) microelectrodes (Utah array, 1mm long and 400 μm apart) and 9 (3 x 3) ECoG electrodes by Ad-Tech Medical Instrument. More details of this custom-made hybrid array, including its implantation, receptive field location, and electrode selection (5 out of 9 ECoG electrodes) are described in our previous study (Monkey 3 of (Dubey & Ray, 2019)). At the time of recording, the microelectrode array had stopped working, so only ECoG data from 5 electrodes were used for analysis. For the second monkey, a 10 x 10 microelectrode array grid (Utah array,1mm long and 400 μm apart) with 96 active platinum electrodes from Blackrock Microsystems was implanted in the primary visual cortex of right cerebral hemisphere (centred at ~12 mm lateral from midline and ~10 mm rostral from occipital ridge). Reference wires were put over the dura near the recording sites. As in our previous studies, only electrodes with reliable and stable receptive field centres across days and with impedances between 250 and 2500 KΩ were used for analysis, yielding 20-21 electrodes. Raw signals were recorded using 128-channel Cerebus Neural Signal Processor (Blackrock Microsystem), bandpass filtered between 0.3 Hz (Butterworth filter, 1^st^ order, analog) and 500 Hz (Butterworth filter, 4^th^ order, and digital), sampled at 2000 Hz and digitized at 16-bit resolution.

### Experimental setup

Visual stimuli were presented using a LCD monitor (BenQ XL2411, 1280 x 720 resolution, 100 Hz refresh rate), gamma corrected and calibrated to a mean luminance of 60 cd/m^2^. Human subjects sat in front of the monitor in a dark place covered by black curtains, with their head movement restricted using a chin rest placed in front of them. Monkeys sat on a primate chair with their head restrained inside a faraday cage to reduce any external electrical noise. Human subjects viewed the monitor from 58 cm, such that the full screen gratings covered the width and height of 46.8° and 27.2° of visual field. Monkeys viewed the monitor from 50 cm, such that the gratings covered 56° x 33° of visual field. Humans and monkeys were required to hold fixation within 2.5° and 2°, respectively, of a small spot of 0.1° in the centre; trials in which fixation was broken were aborted immediately. Eyes were tracked using EyeLink 1000 (sampled at 1000 Hz) for humans and ETL-200 Primate Eye Tracking System (ISCAN, sampled at 200 Hz) for monkeys. In human recordings, each trial began with onset of fixation where they were required to hold their gaze for 2000ms, after which two to three stimuli appeared for 2500, 800 and 1500 ms with interstimulus intervals of 2500, 700 and 1500 ms in the three experiments, respectively. In monkey recordings, each trial started with the onset of fixation where they were required to hold their gaze for 2000 ms, after which two stimuli appeared for 1500 ms with interstimulus intervals of 1500 ms. Monkeys were rewarded with a drop of juice after every successful trial.

### Visual stimuli

In all three experiments, two fully overlapping full screen counter-phase gratings were used to generate a plaid stimulus. Each of the constituent gratings could have a different orientation, contrast, and temporal frequency.

Experiment 1 was aimed to replicate the findings of our previous study (Salelkar and Ray, 2020) in human EEG recordings. Here, the target grating was presented either at an orientation of either 0□ or 90□, contrast of 0 or 50% and temporal frequency of 16 Hz. The mask grating was presented at a range of different orientations (n = 4; 0□, 30□, 60□, 9□), 50% contrast, and 7 different temporal frequencies (10, 12, 14, 16, 18, 20, 22 Hz). When the target grating was absent (0% contrast), the SSVEP response was only due to the mask grating, which allowed us to get the SSVEP gain function (Figure 1A). The spatial frequency of both target and mask grating was fixed at 2 cycles per degree. Total number of stimulus conditions in experiment 1 were 56 (2 target contrasts x 4 mask orientations x 7 mask temporal frequencies). Each stimulus condition on average had about 10 repeats for each subject.

In experiment 2, temporal frequency of target grating was fixed at 15 Hz and temporal frequency of mask grating ranged from 1 to 29 Hz (n = 15; 1, 3, 5, 7, 9, 11, 13, 15, 17, 19, 21, 23, 25, 27 and 29 Hz). We shifted to 15 Hz to minimize sub-harmonic and harmonic responses of target frequency (for 16 Hz, we got weak sub-harmonic responses at 8, 4 and 2 Hz). The contrast, orientation and spatial frequency of target/mask grating were same as experiment 1. Total number of stimulus conditions in experiment 2 were 120 (2 target contrasts x 4 mask orientations x 15 mask temporal frequencies). Each stimulus condition on average had 10 repeats. Due to large number of stimulus conditions, recordings were done in 2 sessions, each lasting two hours on average; stimulus repeats were pooled across the session for analysis.

Experiment 3 was conducted on both human and monkeys. Here, multiple target stimuli were used. To reduce the number of conditions, the number of mask orientations was reduced to 2 (parallel and orthogonal). Target grating was presented at 7 and 15 Hz in human recordings and at 7, 11 and 15 Hz in monkey recordings (recorded in 3 separate sessions, each with one of the three target frequencies). Mask grating was presented at 15 temporal frequencies (1 to 29 Hz, 2 Hz resolution) in human recordings and at 29 temporal frequencies (1 to 29 Hz, 1 Hz resolution) in monkey recordings. The contrast of the target/mask grating was 50% in human recordings and 25% for monkey recordings (to remain consistent with the previous study). Target grating orientation was fixed at 0□, and mask grating was at 0□ and 90□. The spatial frequency of target/mask grating was fixed at 2 cycles per degree. In human recordings, we performed two back-to-back sets of recordings. First, we used one of the target frequencies and presented it at either 0% or 50%, yielding 60 stimulus conditions (2 target contrasts x 2 mask orientations x 15 mask temporal frequencies). Subsequently, we presented the second target frequency at 50% contrast, yielding 30 conditions (1 target contrast x 2 mask orientations x 15 mask temporal frequencies). This is because the 0% target contrast condition which yielded the SSVEP gain function could be estimated from the first set itself. For monkey recordings, we used 1 target frequency in each session, yielding 116 stimulus conditions (2 target contrasts x 2 mask orientations x 29 mask temporal frequencies) per session. Each stimulus condition on average was repeated 10, 10 and 15 times for human, Monkey 1 and Monkey 2, respectively.

### Data analysis

Data were analysed using custom codes written in MATLAB (The MathWorks, Inc). In human recordings, baseline period was defined as −500 ms to 0 (0 being the stimulus onset time) and stimulus period was defined as 250 to 750 ms, yielding a frequency resolution of 2 Hz. In monkey recordings, baseline period was −1000 ms to 0 and stimulus period was 250 to 1250 ms, yielding a frequency resolution of 1 Hz.

EEG data was subjected to a series of processing pipelines before using it for further analysis. First, bad stimulus repeats were removed using the pipeline described in our previous paper (Artifact rejection of (Murty et al., 2020)), based on which almost ~20% of stimulus repeats were rejected. In this pipeline, any stimulus repeat in which either the raw waveform or power spectral density (PSD) was more than 6 standard deviation away from their mean across all repeats was tentatively labelled ‘bad’. Next, electrodes with more than 30% of all repeats labelled as ‘bad’ were marked as bad electrodes and removed from analysis. A stimulus repeat was then marked ‘bad’ when it was present in more 10% of the remaining electrodes. Finally, any electrode for which the mean PSD had a slope less than zero between 56-84 Hz in the baseline period (indicating a flat PSD) was also discarded. This procedure yielded a set of good electrodes for each subject, and a set of good stimulus repeats that were common for all good electrodes.

Second, subjects or sessions with less than 2 repeats for each stimulus condition were also removed. Third, we computed SSVEP amplitude spectra by taking the Fast Fourier Transform (FFT) of averaged signal across all the repeats deemed good for a stimulus condition, and rejected electrodes with target SSVEP responses of less than 0.5 μV in EEG and 2.5 μV in ECoG and LFP. Finally, human subjects with less than 2 good electrodes (out of 8) were rejected. Overall, this yielded 8 (out of 10), 8 (out of 10) and 7 (out of 11) good subjects for the three experiments, respectively. In monkey recordings (experiment 3), we obtained usable data from 2 to 5 ECoG electrodes in the three sessions (one for each target frequency) from Monkey 1 and 20-21 LFP electrodes from Monkey 2 in all sessions. Counterphasing stimuli used in this study produces a prominent response at twice the stimulus frequency, since neurons respond to luminance changes in either the positive or negative direction (two responses per stimulus cycle). The amplitude difference was therefore calculated as *Amp_ST_*(2 * *f*) – *Amp_BL_*(2 * *f*), where *Amp_ST_*(2 * *f*) and *Amp_BL_*(2 * *f*) are the amplitudes at twice the target/mask frequency in stimulus and baseline period respectively.

### Statistical analysis

One-sample t-test was performed to quantify the significance level of differences in high and low frequency responses.

## Results

### Asymmetric suppression shown previously in monkey recordings is also present in human EEG

We first tested whether asymmetric suppression observed in macaque LFP responses (Salelkar and Ray, 2020) is also present in human EEG. We recorded EEG from parietooccipital and occipital electrodes of 8 healthy young adults (5 females). As in our previous study, we presented two overlapping gratings (i.e., a plaid stimulus), with both constituent gratings counter-phasing at different frequencies. One grating, which we call the “target,” always counter-phased at 16 Hz, and produced a strong SSVEP response at 32 Hz. The orientation of this target grating was either 0° (5 out of 10 subjects) or 90° (remaining subjects). The second (“mask”) gratings had variable temporal frequency (n =7; 10, 12, 14, 16, 18, 20, 22 Hz) and orientation (n = 4; 0°, 30°, 60°, 90°). The contrast of the mask frequency was fixed at 50%, while the target grating was either 0% or 50%.

Figure 2A shows the amplitude spectra of EEG responses of one subject, averaged over all electrodes that qualified the thresholding criteria (N = 8; see Methods for details). The response of the plaid was evaluated as the change in amplitude between grating-only and plaid conditions (blue and color traces) at 32 Hz (Figure 2B). These were generally negative, indicating target frequency SSVEP suppression by the mask. Positive responses were observed when both gratings had the same temporal frequency. In the parallel condition, we observed substantially stronger suppression of amplitude response of plaids with mask frequencies less than 16 Hz target (e.g., compare 10 Hz mask frequency; row 1) versus more (22 Hz; row 7). We quantified the asymmetry by subtracting the average response of plaids with mask frequencies greater than the target (18, 20 and 22) from the lower ones (10, 12 and 14), which showed stronger asymmetry in the parallel condition compared to orthogonal conditions (Figure 2C), although the difference was significant at all orientations (N=8, p-value = 0.0042 (parallel), 0.0007 (30° separation), 0.0001 (60° separation), 0.0126 (orthogonal)).

**Figure 2:**
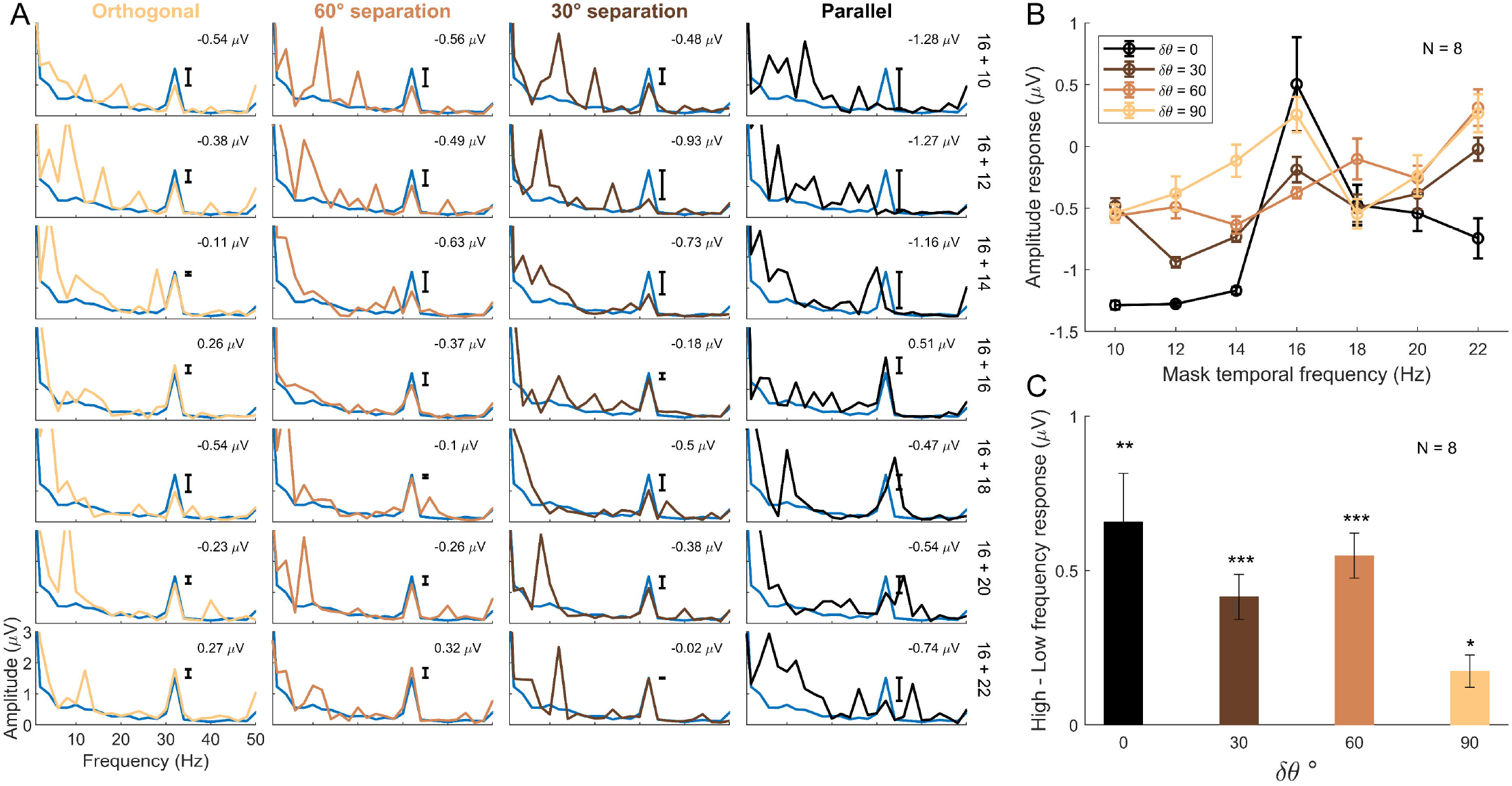
EEG amplitude response suppression of Subject 1. **(A)** The amplitude spectra are averaged over the good electrodes (N = 8). Each row indicates a different temporal frequency of mask grating (indicated at extreme right of the fourth column) that is superimposed with a grating with temporal frequency of 16 Hz (the target grating); the columns show different orientations of mask grating relative to the target grating. Blue trace is the 16 Hz “grating-only” condition (same trace in all plots). The number in the inset is the difference in amplitude at the target frequency (i.e., 32 Hz) between the grating-only condition and the plaid condition (i.e., between the blue and color traces at 32 Hz). **(B)** Difference between the grating-only condition and plaid condition as described above, plotted as function of mask frequency for different orientations. **(C)** Average difference between amplitudes above (18, 20 and 22 Hz) and below (10, 12 and 14 Hz) the target frequency (16 Hz), for different orientation difference conditions. The error bars indicate the standard error of mean along with the significance level (***: p < 0.001; **: p < 0.01; *: p < 0.05).

Results were consistent for averaged response across all subjects (N = 8; Figure 3A and B). Because these conditions were also tested in experiment 2 (except for a shift in the target frequency from 16 Hz to 15 Hz), we combined the subjects from the two experiments (N = 16; Figure 3C). Asymmetry in suppression decreased with orientation dissimilarity, although remained significant in all conditions (p-value = 0.0011 (parallel), 0.0021 (30° separation), 0.0001 (60° separation), 0.0055 (orthogonal)).. These results are consistent with our previous study (Salelkar & Ray, 2020), and consistent with both the gain-specific and low-frequency specific hypotheses (Figure 1D and 1E).

**Figure 3:**
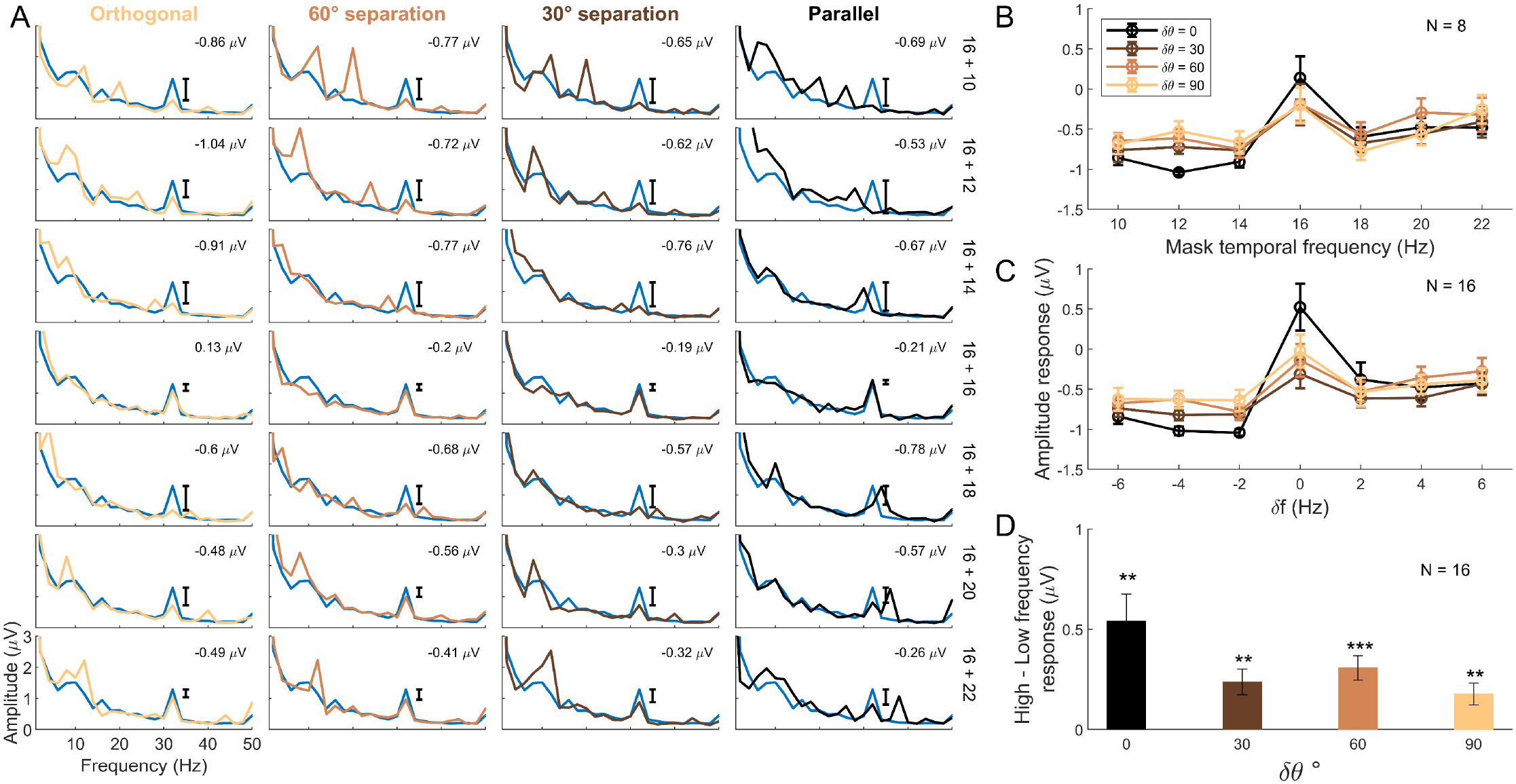
Averaged EEG amplitude response suppression of all subjects. **(A-B)** Same as 2A-B, but for data averaged over all usable subjects (N = 8) **(C)** Same as B, but after adding data from 8 subjects who participated in experiment 2. Amplitude response is plotted as a function of delta frequency (mask frequency – target frequency), since for the second experiment the target frequency was 15 Hz. **(D)** Same as Figure 2C, but for data averaged across 16 subjects.

### Target frequency suppression is not low-frequency tuned

In our previous study, we dissociated the two hypotheses by presenting the target at a different frequency (8 Hz). Here, we tested a different prediction. While keeping the target frequency fixed, if we increase the range of mask frequencies, the two hypotheses predict different behaviour at low mask frequencies (compare the magenta traces in Figure 1D versus 1E). In experiment 2, we presented two overlapping counter-phase gratings with target grating at 15 Hz at 0□ (5 out of 10 subjects) or 90□ (remaining subjects) orientation, and varied the temporal frequency (n =15; 1 to 29 Hz in steps of 2 Hz) and orientation (n = 4; 0□, 30□, 60□, 90□) of the mask grating. Human EEG was recorded from parieto-occipital and occipital electrodes of 8 (5 females) healthy young adults. The target frequency was changed from 16 to 15 Hz only to avoid weak sub-harmonics at 8, 4 and 2 Hz when the target was at 16 Hz, since we were interested at the low-frequency masking frequencies.

Figure 4 depicts the averaged amplitude spectra of 8 subjects (same format as Figure 2 and 3). In the orthogonal condition, symmetric suppression was observed (non-specific hypothesis; Figure 1C). Surprisingly, in the parallel condition, the suppression profile was more similar to SSVEP-gain specific hypothesis (Figure 1D) than low-frequency specific (Figure 1E), inconsistent with our previous results (Salelkar and Ray, 2020). To quantify these results, we computed the difference between the mean amplitude response for mask frequencies 1, 3 and 5 Hz and the mean amplitude response for mask frequencies 9, 11 and 13 Hz (these data points are shown as asterisks in Figure 1 and as thick filled circles in 4B). This quantity is expected to be near zero for non-specific suppression, negative for SSVEP-gain specific hypothesis and positive for low-frequency hypothesis (Figure 1B, magenta bars). The difference was significantly negative for parallel and 30° orientation difference (p-value = 0.0154 (parallel), 0.0194 (30° separation), 0.0740 (60° separation), 0.3522 (orthogonal)), consistent with the SSVEP gain-specific hypothesis.

**Figure 4:**
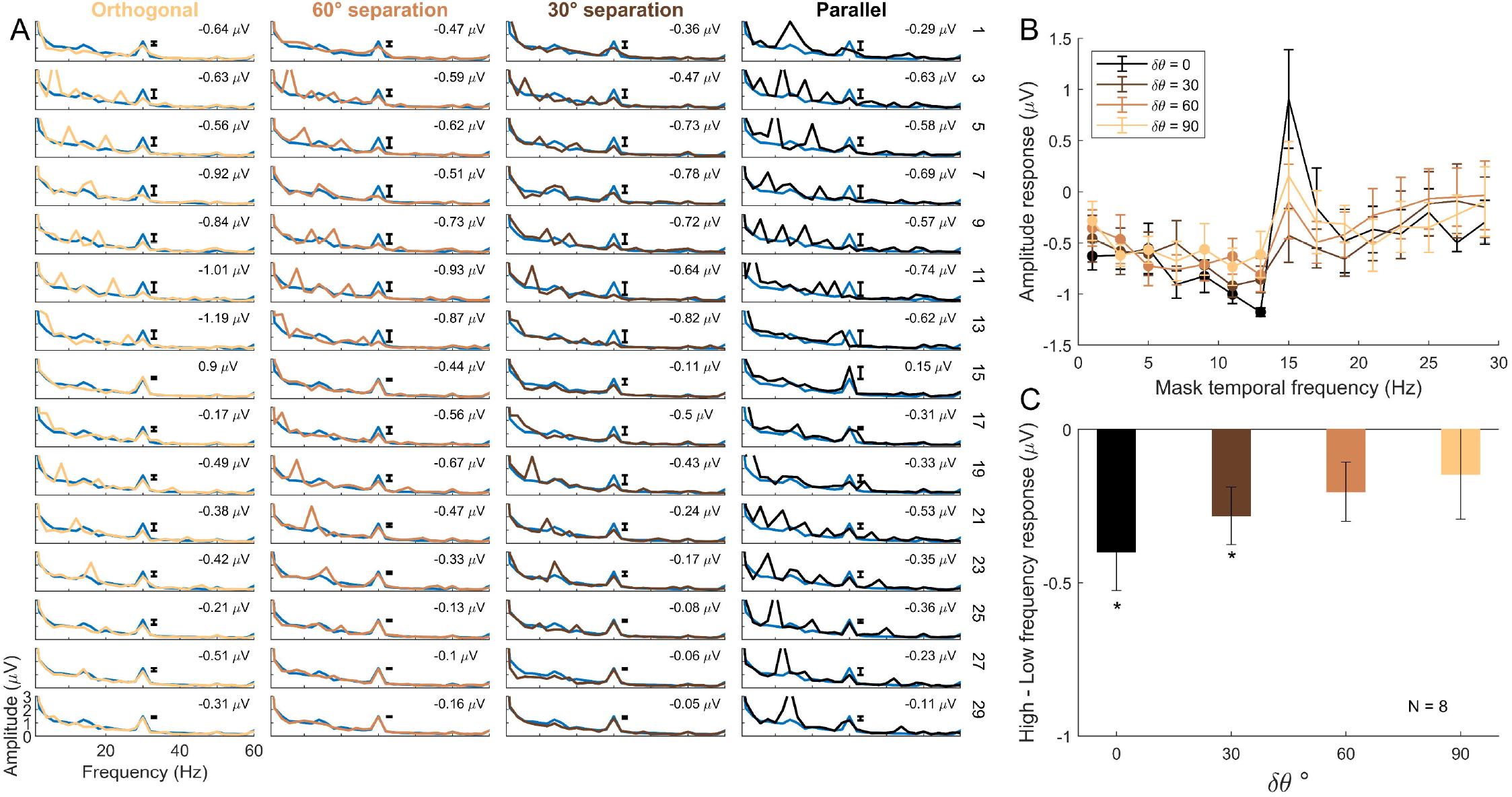
Averaged EEG amplitude response suppression of all subjects for a wider range of mask frequency. **(A-B)** Same format as in Figure 3A-B, but for 8 subjects who participated in experiment 2 which had a wider range of mask frequencies and a target frequency of 15 Hz. **(C)** The average difference between responses obtained for plaids with mask frequencies of 9, 11 and 13 Hz and mask frequencies of 1, 3 and 5 Hz (as indicates in solid circles; Figure 1B shows this value under different hypotheses).

### Suppression profile is dependent on the target frequency

The three hypotheses described in Figure 1C-E all have a “global” suppressive function, as shown in green traces in Figure 1. Another hypothesis, not tested previously, is that the suppression also has a “local” component, arising out of local interactions between target and mask frequencies. This would lead to different suppression profiles dependent on the target frequency, as shown in Figure 1F, and could reconcile our previous results.

We tested this prediction in both monkey and human recordings, including one monkey who was part of the previous study (Monkey 1; only ECoG data could be recorded because the microelectrode array had stopped working). We used two overlapping full screen counterphase gratings with target frequencies set to either 7, 11 or 15 Hz (7 and 15 Hz for humans) and mask frequencies that varied from 1 to 29 Hz (n = 29 for monkeys, n=15 for humans; see Materials and Methods for more details).

Figure 5A shows the amplitude response as a function of mask frequency for the two monkeys (columns 1 and 2) and human EEG, in the parallel condition. Consistent with the new hypothesis, the suppression profile changed systematically depending on the target frequency. The difference between the responses from plaids with mask frequencies higher versus lower than 7 Hz (i.e, average between 8-13 Hz and 1-6 Hz, shown in solid circles) was positive when the target was at 7 Hz (Figure 5C, blue traces), consistent with our previous report (Salelkar and Ray, 2020), but became progressively negative as the target frequency shifted to 15 Hz. This effect was largely abolished in the orthogonal condition (Figure 5B, C).

**Figure 5:**
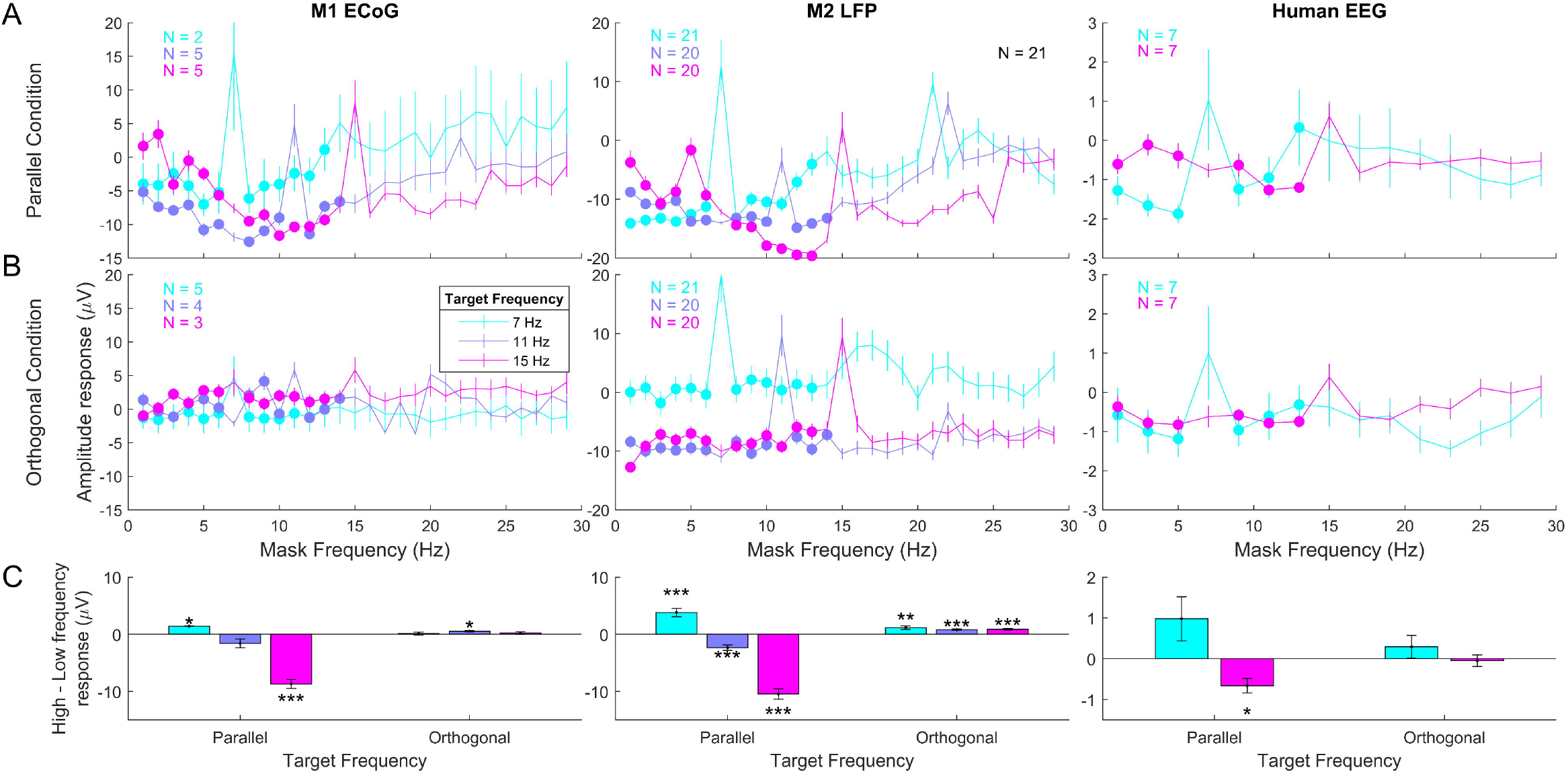
Suppression profile for different target frequencies. **(A)** SSVEP amplitude suppression for target frequencies 7, 11 and 15 Hz in monkey recordings and 7 and 15 Hz in human recordings as a function of mask temporal frequency in the parallel condition. **(B)** Same as A, but for the orthogonal condition. **(C)** For 7 and 15 Hz target frequencies, the average difference between responses obtained for plaids with mask frequencies of 8 to 13 Hz and mask frequencies of 1 to 6 Hz in monkey recordings. In human recordings, average difference between responses obtained for plaids with mask frequencies of 9, 11 and 13 Hz and mask frequencies of 1, 3 and 5 Hz. For 11 Hz target frequency in monkey recordings, the average difference between responses obtained for plaids with mask frequencies of 8 to 14 Hz (except at 11 Hz) and with mask frequencies of 1 to 6 Hz (as indicated in solid circles).

## Discussion

We used two overlapping counter-phasing gratings at varying relative orientations and temporal frequencies to comprehensively map the interactions between the two SSVEPs in human EEG. We confirmed that the low-frequency suppression (in which lower temporal frequencies cause larger suppression) for parallel but not orthogonal gratings as described recently in monkey LFPs (Salelkar & Ray, 2020) was also observed in human EEG. However, this low-frequency suppression gradually diminished as the difference between the target and mask frequency increased, inconsistent with the hypothesis proposed earlier (Salelkar & Ray, 2020). Instead, we found that the overall suppression profile was dependent on the target frequency in both human EEG and monkey LFP/ECoG data, which reconciled previous results. Together, these results provide a comprehensive account of the suppressive effects of two SSVEP tags along temporal frequency and relative orientation domains.

Previous psychophysical studies measuring contrast sensitivity as a function of mask frequency provide important clues about the mechanisms that can potentially explain our results. These studies have reported the presence of at least two temporal frequency masking channels in the visual system: lowpass and bandpass (Anderson & Burr, 1985; Boynton & Foley, 1999; Cass & Alais, 2006a; Hess & Snowden, 1992). The frequency cut-off of lowpass masking channel was observed to be 10 Hz. The bandpass masking channel response was observed in the frequency range of 7 to 13 Hz with peak around 10 Hz which closely resembled our SSVEP response function or the global profile (Figure 1A). The two channels might interact with each other and engage different inhibitory mechanisms, depending on the target frequency. Low and high target frequency have shown to involve lowpass and bandpass masking mechanisms respectively (Anderson & Burr, 1985; Cass & Alais, 2006a).

In our experiments with parallel masks, for target frequency of 15 Hz, the suppression profile was indeed bandpass and similar to the global profile, while it was low-pass for 7 Hz target frequency, consistent with the idea that different masking mechanisms were involved at different target frequencies. However, not all results in these psychophysics studies were consistent with our results. For example, some studies have suggested asymmetric suppression where high mask frequency suppressed low target frequency but not vice-versa (Allison et al., 2001; Cass & Alais, 2006a), but we observed asymmetry in the opposite direction. It should be noted that there are important differences between the stimuli used: we used counter-phasing gratings to generate SSVEPs, while many of these previous studies used drifting gratings (a counterphase grating can be decomposed into two drifting gratings moving in opposite directions). Further, these studies measured behaviour (for example, masking threshold), as opposed to a neural measure (SSVEP amplitude) reported here. If some of these temporal frequency channels have non-cortical origins, then the neural responses from a cortical area may differ from behavior. For example, Cass and Alais (2006) suggested that band-pass channel is pre-cortical while the low-frequency channel has cortical origins.

In previous studies with EEG recordings, Regan (1983) studied SSVEP responses while presenting target and mask gratings at varying spatial and temporal frequencies. The target response was heavily suppressed by masks that had similar spatial frequency as the target, indicating the presence of multiple narrow bandwidth spatial-frequency tuned mechanisms, consistent with psychophysics as well (eg. Anderson and Burr, 1985). For temporal frequency, the interaction (recorded from a single human subject) was consistent with our results, so our findings generalize these results to a larger pool of subjects and over multiple scales of electrophysiological signals. In addition, that study did not report the effect of relative orientation of the two gratings. A SSVEP study that looked at the effect of orthogonal versus parallel masks found stronger suppression with orthogonal masks, contrary to our results (Burr & Morrone, 1987). However, this study used a variety of different spatial frequencies and contrasts as well, while we used a fixed spatial frequency (2 cycles/degree) and contrast (50%). Given the dependence of masking strength on such parameters, it is difficult to directly compare these results.

More recently, masking phenomena have been explained using normalization, a neural computation observed throughout the brain describing response modulation due to multiple stimuli (Carandini & Heeger, 2012). According to the model, the excitatory drive due to one stimulus is normalized by inputs from the surrounding neuronal population having broader tuning properties (Carandini et al., 1997; Carandini & Heeger, 1994; Heeger, 1992). The normalization model initially proposed for single unit recordings has also been used to explain temporal frequency masking effects in human EEG studies (Boynton & Foley, 1999; Candy et al., 2001; Tsai et al., 2012). These models could represent target, mask as well as intermodulation (IM) response profiles as a function of target/mask contrasts by manipulations in the way excitatory and inhibitory inputs were summed and passed through simple non-linearities (such as rectification or squaring). We also recently used a tuned normalization model to capture the signal contributing to target SSVEP response suppression as a function of mask frequency (Salelkar & Ray, 2020). The low-frequency suppression was modelled by essentially assuming a particular type of suppression function (green trace in Figure 1C-E, which was assumed to be a downward sloping linear function; see (Salelkar & Ray, 2020) for details).

In the previous paper, we discussed a potential reason that could lead to such a low-frequency suppression profile (Figure 1E), and here we further speculate how some changes in this model can explain the local interactions between target and mask gratings as reported in this paper (Figure 1F). To explain the low-frequency suppression profile, we first consider a variant of the normalization model proposed earlier (Baker & Wade, 2017; Foley, 1994). In these models, two input sinusoids each for target and mask frequency (sin ω_T_t and sin ω_M_t) act as the excitatory drive which is the numerator of the response. The denominator (normalization) part of the model is the inhibitory drive which is the combined responses of target and mask frequency after passing through some non-linearity. For example, consider the response of the form: R= (sinω_T_t+ sinω_M_t)^2^/[sin(ω_T_t)^2^ + sin(ω_M_t)^2^ + σ]. By adding a squaring non-linearity in the numerator, the response has strong responses not only at 2ω_T_ and 2ω_M_ but also at the intermodulation terms |ω_T_ + ω_M_| and |ω_T_ – ω_M_|, as observed in data (Figures 2–4; Foley, 1994). Further, by changing whether the non-linearity is applied before or after summation, or the exponent of the non-linearity, the behaviour of the model can be modified.

Simulating the response as per the model described above immediately presents a big problem: the presence of the rapidly varying sinusoidal signals in the denominator lead to spurious peaks at many frequencies beyond ω_T_, ω_M_ and their harmonics and sum/differences, which is never observed in real data. This issue can be addressed by assuming that the normalization signal varies slowly over time, which can be achieved by low-pass filtering the normalization signal (Tsai et al., 2012). However, this procedure also makes the normalization strength stronger when the mask frequency is low, in effect generating a low-frequency suppression profile as shown in Figure 1E. A simple way to achieve target frequency specific suppression (Figure 1F) in this framework is to sum the sinusoids before squaring and low-pass-filtering in the normalization term, i.e. by assuming that the normalization signal is low-pass-filter{(sinω_T_t + sinω_M_t)^2^ + σ}. This is because now expanding the sinusoidal terms produces a |ω_T_ – ω_M_| IM component, which has very low frequency when target and mask frequencies are close to each other and therefore is preserved after low-pass-filtering, while other frequencies are filtered out. This model explains the strong suppression of target SSVEP response when mask frequency is in vicinity of the target frequency, leading to different suppression profiles depending on the position of the target frequency (Figure 1F). Further, the observed asymmetry around the target – where masks below the target frequency produce stronger suppression, can also be explained because slower mask frequencies in general get less attenuated by the low-pass filter compared to higher mask frequencies (this is the original low-frequency suppression argument discussed earlier).

Although in our simulations with this model we were able to reproduce the target-specific suppression, we have refrained from developing a formal model here because the model is highly unconstrained (similar to our previous results in Salelkar and Ray, 2020). The model parameters are fitted as each target frequency separately (see Salelkar and Ray, 2020 for details), so to adequately constrain such models, multiple contrast levels of mask and target frequencies are needed for each combination (as done, for example, in Foley 1994 or Anderson and Foley, 1999). In our experiments we used only one contrast level of target and mask gratings due to large number of stimulus conditions that led to long recording hours (~3 hours for Experiment 3). We noticed that human SSVEP responses sometimes showed minor variations when recorded on different days, and therefore performed all recordings in a single day. Along with multiple contrast levels for each target and mask frequency, multiple spatial frequencies also need to be tested to fully characterize the interactions between two flickering stimuli, for which monkey recordings with chronically implanted arrays may be the ideal recording platform. It would also be interesting to see how these results relate to the spiking unit activity. In the current monkey recordings, we used full screen flickering stimuli to be comparable to the human EEG recordings, which led to a small number of stable spiking units in Monkey 2.

Apart from classic attention studies, SSVEP paradigms have become popular in Brain-Computer Interface (BCI) applications, where two or more simultaneously presented stimuli are tagged with different flickering frequencies, and relative change in SSVEP response when one stimulus is attended versus unattended is used to control an application (Ding et al., 2006). A better understanding of the interactions between multiple flickering stimuli at a neural level holds promises towards improvement in such BCI applications, and also enhances our understanding of how the visual system represents such competing stimuli.

## Acknowledgements

This work was supported by Department of Biotechnology (DBT)/ Wellcome Trust India Alliance (IA/S/18/2/504003; Senior fellowship to SR) and the DBT-IISc Partnership Programme.

